# Forecasting of a complex microbial community using meta-omics

**DOI:** 10.1101/2022.10.19.512887

**Authors:** F. Delogu, B. J. Kunath, P. M. Queirós, R. Halder, L. A. Lebrun, P. B. Pope, P. May, S. Widder, E. E. L. Muller, P. Wilmes

## Abstract

Microbial communities are complex assemblages whose dynamics are shaped by abiotic and biotic factors. A major challenge concerns correctly forecasting the community behaviour in the future. In this context, communities in biological wastewater treatment plants (BWWTPs) represent excellent model systems, because forecasting them is required to ultimately control and operate the plants in a sustainable manner. Here, we forecast the microbial community from the water-air interface of the anaerobic tank of a BWWTP via longitudinal meta-omics (metagenomics, metatranscriptomics and metaproteomics) data covering 14 months at weekly intervals. We extracted all the available time-dependent information, summarised it in 17 temporal signals (explaining 91.1% of the temporal variance) and linked them over time to rebuild the sequence of ecological phenomena behind the community dynamics. We forecasted the signals over the following five years and tested the predictions with 21 extra samples. We were able to correctly forecast five signals accounting for 22.5% of the time-dependent information in the system and generate mechanistic predictions on the ecological events in the community (e.g. a predation cycle involving bacteria, viruses and amoebas). Through the forecasting of the 17 signals and the environmental variables readings we reconstructed the gene abundance and expression for the following 5 years, showing a nearly perfect trend prediction (coefficient of determination ≥ 0.97) for the first 2 years. The study demonstrates the maturity of microbial ecology to forecast composition and gene expression of open microbial ecosystems using year-spanning interactions between community cycles and environmental parameters.

## Introduction

Microorganisms are ubiquitous on planet Earth^1^ and constitute up to 17% of its carbon biomass^2^. Microbial lineages are continuously evolving to fill a diverse set of ecological niches, balancing their complementary metabolic capabilities to form communities^1^, which, in turn, affect biogeochemical cycles^3^. Understanding the temporal dynamics of microbial ecosystems and their links to the environment has become a common problem for many research fields spanning biomedicine, agriculture, biotechnology and climate change. Whilst forecasting community composition dynamics has been successfully achieved for some environments (e.g. Larsen et al.^4^ and García-Jiménez et al.^5^), the forecasting of their gene expression dynamics over time and environmental conditions remains an open challenge^6^. One reason for this is the lack of a generalised framework that enables us to capitalise on the recent advances in the meta-omics field. The surface community of Biological Wastewater Treatment Plants (BWWTPs) represents an excellent candidate to become a model system to study the forecasting of microbial behaviour (dynamics of the populations and their related gene expression) for a threefold reason^7^. Firstly, BWWTPs share the challenges linked to most environments, as it is an open system, with a constant influx of new populations^8^ and exchange of matter and energy with the environment (i.e. access to open air and sun irradiation). However, these challenges can be mitigated by keeping operational parameters (e.g. pH, phosphate and nitrate) within a controllable range. Secondly, BWWTPs communities share common metabolic pathways, albeit every local community has its own equilibrium and its detailed makeup depends on the operational parameters, geographical location and inflow composition^9^–^11^. Microbial communities in WWTPs possess dynamics at different temporal scales that are rather well described: the microbial and chemical composition of the inflow is known to change according to the time of the day, the day of the week and the inflow volume^12^. In addition, temperature-driven seasonality has been found to influence the community^9,13^ as well as multi-annual trends^14^. While one-time destructive perturbations show an impact on the community such as human intervention (e.g. bleaching, shutdowns)^14,15^ and weather (i.e. rain)^16^, they are all monitored or encoded in the standard operational parameters of the plants. Finally, forecasting the behaviour of microbial communities in BWWTPs is highly desirable as stable operation allows reclamation of clean water as well as the harnessing of chemical energy^17^ while at the same time its functioning has to minimise the production of the greenhouse gases such as N_2_O^18^.

There exist several categories of time-series analysis. These are based on: i) previous knowledge (such as curve fitting^19^ and classification^20,21^), ii) subsetting (e.g. segmentation^22^), iii) clustering (e.g. based on various metrics such as Euclidean Distance^23^ or Dynamic Time Warping^24^) iv) prediction (such as forecasting^25^ and intervention analysis^26^), and v) decomposition (e.g. Singular Value Decomposition - SVD^27^). The prediction of future states of ecological communities and their interplay with the environment have been successfully tackled in the case of available interaction models and/or limited number of species^28,29^. However, predictions of microbial metabolic behaviour are rendered challenging for naturally occurring microbial ecosystems as well as industrially-relevant ones, such as in BWWTPs. In this context, metagenomics (MG)^30,31^, metatranscriptomics (MT)^32^ and metaproteomics (MP)^33^ enable the establishment of sample-specific reference databases that simultaneously resolve both compositional and functional aspects of the system. When dealing with complex and uncharacterised microbial systems, far from lab-scale experiments, empirical modelling can enable efficient representation and forecasting. To achieve this we foresee a combination of strategies to extract all the temporal information in an agnostic manner, such as through SVD, and to perform forecasting by explicitly computing the temporal cycles and link those patterns directly to the explanatory variables. SVD can decompose a matrix in two separated matrices of eigenvectors and a vector of eigenvalues. When applied to gene abundance (or expression) data over time, one of the matrices is interpreted as the set of temporal patterns underlying the data and the other as the “loadings” (i.e. how much each individual gene is contributing to each pattern). The seasonal version of the forecasting method AutoRegressive Integrated Moving Average (ARIMA) computes cyclical (seasonality), autoregressive (temporal self-dependence), differencing (difference between consecutive time points) and moving-average (averaging of consecutive time points) components of a time-series^34^. It thereby offers a very flexible framework for time-series modelling^34^.

We present a general analytical framework which is capable of exploiting the richness of temporal multi-layered meta-omics data in the context of microbial communities. We demonstrate its power through the analysis of a Lipid Accumulating Organisms (LAO) surface community (LAO) from an anaerobic tank of the BWWTP in Schifflange (Luxembourg). The sampling spans more than one year with 51 samples collected from March 2011 to May 2012 from which we co-extracted the macrobiomolecules and analysed the derived MG, MT and MP datasets^35^ alongside the physicochemical factors measured at the site. We reconstructed the MG structure of the community, alongside its taxonomy, genetic potential, transcript and protein levels. We employed SVD to extract relevant temporal patterns, which were then clustered into 17 fundamental signals. Those were integrated with collected environmental parameters to build an ARIMA model, augmented with seasonal components, which could explain the observed signals. Multiple models (ARIMA, prophet^36^ and NNETAR neural networks model^37^) were trained to forecast the following five years’ signals. Validation was conducted using future time points, i.e. 21 samples covering the months of June for the years 2012-2016. This allowed us to correctly predict the gene abundance and expression of the populations in the community.

## Results and Discussion

### Functional and genetic characterization of LAO

From the experimental period between 2011-03-21 and 2012-05-03^38^ we obtained 51 weekly samples, that were submitted to multi-omic analyses (MG, MT and MP) and analysed individually to obtain 51 genomic assemblies, collections of metagenome-assembled genomes (MAGs), plasmids, viruses, unbinned prokaryotic chromosomal contigs and the corresponding gene expression at the transcriptional and proteomic levels. A week is the In order to form coherent sets spanning the whole time-series, we individually clustered the bins (prokaryotic and eukaryotic) and the contigs (viral, plasmid and unbinned) according to their sequence (see **methods**), which led to a total of 144 representative MAGs (rMAGs) and 1,681,736, representative contigs (rContigs), yielding 4,711,952 Open Reading Frames (ORFs) (**Supplementary Table 1**). A KEGG Orthology group (KO term) was assigned to 55% of the total retrieved ORFs, whilst taxonomic affiliations were assigned to 38.5%. The number of ORF copies as well as their detected gene expression and protein abundances were determined over the extended dataset (see **methods**). We found on average 2.2×10^6^ (s.d. 4.8×10^5^) ORFs, 9.1×10^5^ (s.d. 1.7×10^5^) transcripts and 2.4×10^5^ (s.d. 2.5×10^4^) protein groups per sample. However, the vast majority of the genes were not found to be expressed over the entire dataset or were only detected in a few samples, with a maximum of 16.8×10^6^ ORFs detected in one sample. This suggests that a significant fraction of the gene pool in LAO is not specifically required for community function but rather their cumulative functional effort may be compartmentalised, fitting the previous results from Roume et al.^39^ showing how a large portion of the community is redundant, and only few functions are keystone. Read recruitment (on the ORFs) per sample was on average 59% (s.d. 9%) for the MG, 82% (s.d. 3%) for the MT, whilst the peptide recruitment was 27% (s.d. 4%).

The rMAGs spanned the expected phyla of the BWWTP community, and included member of the Actinobacteria, Bacteroidetes, Chlorobi, Fusobacteria, Nitrospirae, Proteobacteria and Spirochetes with the addition of the *Candidatus* Gracilibacteria (**Figure 1a**). On a more detailed taxonomic level, we were able to identify three strains of *M. parvicella* and 17 strains of *Moraxella* spp. At no point over the course of the time series did a single rMAG largely dominate the community, but the combined populations of the genera *Microthrix* and *Moraxella* exhibited a percentage abundance with medians of 15.9% and 3.6%, respectively^38^. The majority of the contigs were not affiliated to defined MAGs (**Figure 1b**), and are likely coming from incomplete genomes and alternative regions of the rMAGs, thus encapsulating the within-population diversity of the LAO community.

**Figure 1.**
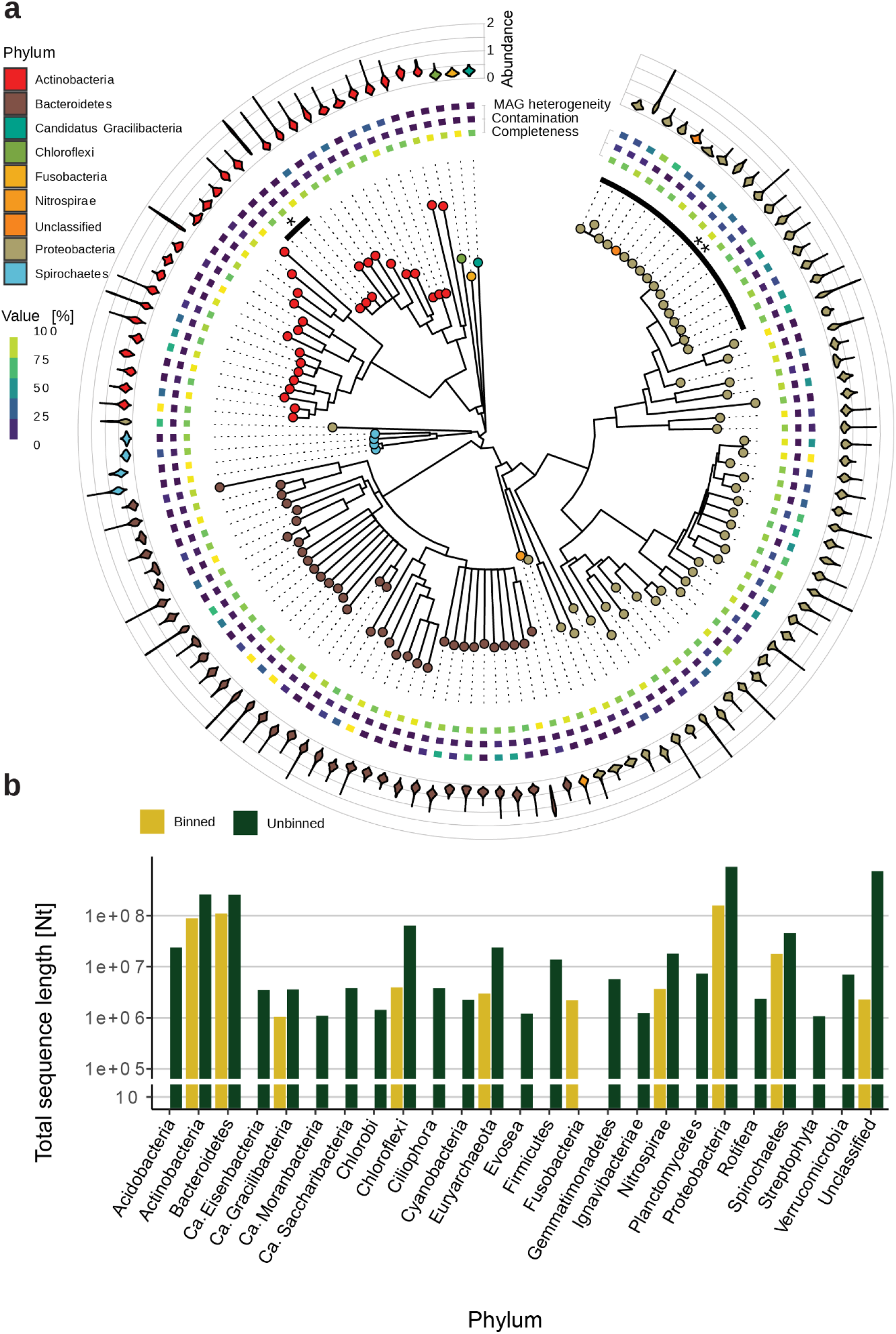
Diversity and quality of the rMAGs and their representativeness in the meta-omics dataset. **a.** The phylogenetic tree of the rMAGs in LAO (generated using gtdb-tk^40^) contains the 125 bacterial rMAGs in the system. The heatmap ring contains the CheckM quality measures per rMAG (completeness, contamination and MAG -originally strain-heterogeneity), which were filtered to be at least 75% complete and at a maximum 25% contaminated (median: 2%). The violin plots contain the time-averaged (train time series) depth profiles over the contigs forming the rMAG. The two sections of the tree noted as * and ** highlight the strains of *M. parvicella* and *Moraxella* sp., respectively. **b.** The cumulative length of the contigs (longer than 1000nt, see **Methods**) for the 25 most abundant phyla displayed for the rMAGs and unbinned contigs.

### The temporal signals underlying the microbial community

Considering that the information necessary to forecast the community dynamics and linked gene expression may be most represented in any biological (e.g. taxonomical or functional representation) or environmental data layer, we decided to include multiple layers in our analysis. Regarding the microbial community we explored multiple taxonomic and functional levels at once and summarised their temporal characteristics. Thus, the three quantification matrices (MG, MT and MP) were used to compute “summary” matrices according to the ORFs’ descriptors. Hence we computed one matrix per omic layer for the six formed taxonomic descriptors (Phylum, Class, Order, Family, Genus and Species) and two functional ones (KO terms and pathways). The resulting 27 matrices (3 original and 24 summary) were used to compute the system’s eigengenes (EGs) i.e. the independent patterns underlying the data and their potential time dependency^27^. In previous works, the first EG in a time-series has been demonstrated to represent the “steady state” gene expression, encapsulating the largest explained variance (EV), and was removed to perform the timeseries analysis^27^. Indeed, the first EG showed the largest variance explained (around 15% in all the datasets), therefore we excluded it from the subsequent analysis. We screened the subsequent EGs for time-dependency (see **Methods**) selecting a set of 210 EGs and assessed how much of the data variation they explained beyond the first EG (**Supplementary Figure 4**).

Considering that the EGs of each matrix are linearly independent (i.e. they do not have redundant information) for each matrix, we expected some level of redundancy by using different types and levels of summarising the information. In order to reduce this redundancy and bring together the same temporal behaviours, we clustered the set of 210 EGs into 17 representative EGs (see **Methods**). These are hereafter referred to as signals (S1-17) and shown in **Figure 2a**. We assumed that the 17 signals were not redundant because they were different enough to not cluster together. Each cluster contained multiple EGs with their associated EV (**Supplementary Figure 5**) and we associated the maximum EV of each cluster to its respective signal. In total, signals S1-17 accounted for 91.1% of the EV in the system (whilst the leftover 8.9% represented noise) and covered all temporal information in the training set.

**Figure 2.**
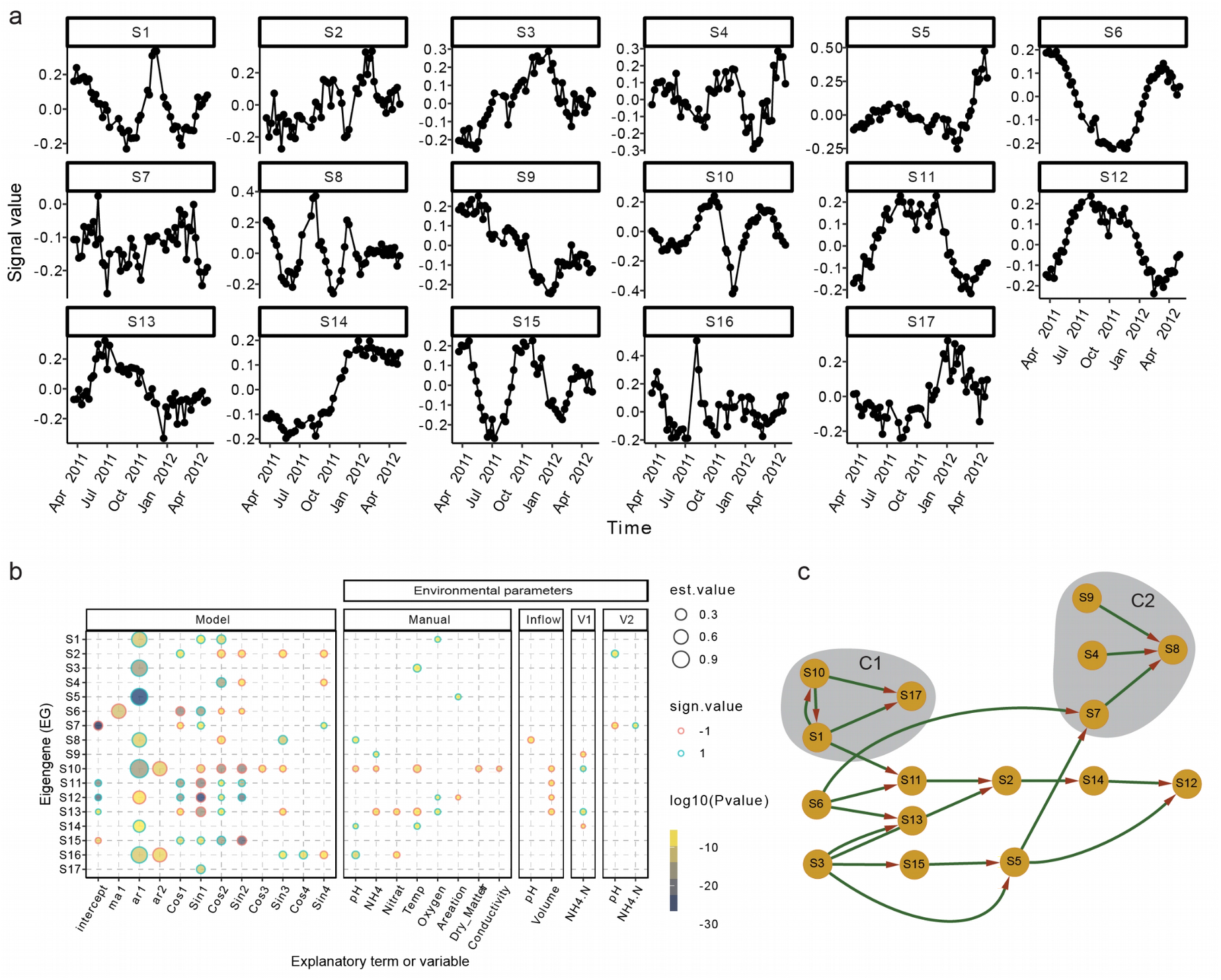
Eigengene modelling using ARIMA augmented with environmental parameters and Fourier terms. **a.** The signals S1-17 encapsulate the time-dependent dynamics underlying the microbial community. The scale of the y-axis is dimensionless as the eigenvectors. **b.** The S1-17 are explained as ARIMA processes under the influence of the environmental variables. The five blocks of explanatory variables are: Model (ARIMA components), Manual (manually collected environmental variables, directly on the sampling location), Inflow (inflow stream of wastewater in the plant), V1 (first anaerobic tank in the plant), and V2 (second anaerobic tank in the plant). Every circle represents a significant variable (Benjamini-Hochberg adjusted p<0.05) for the corresponding signals among S1-17, the size represents the value of the coefficient, the ring colour its sign and the fill colour the log10(p-value). **c.** The signals are connected by a temporal transfer of information, suggesting a succession of ecological events.

The 17 representative signals (S1-17) were modelled using all the non-collinear environmental parameters as variables (see **methods**) as shown in **Figure 2b**. Moreover, the model includes predictors derived from the ARIMA, such as the intercept (the basal abundance/expression), autoregression (the time-lagged self dependence) and sine/cosine (the cyclical behaviours, including seasonality), which explain the microbial process through its mathematical components. In summary, we include self-dependent, cyclical and environmental interactions to explain community dynamics. As seen in **Figure 2b**, all the signals are generally explained more via the mathematical variables rather than the environmental ones. This is partially due to the fact that some of the environmental variables have a seasonal trend too (e.g. temperature) and their impact will be significant in the model if their values explain more than the seasonality (i.e. having a fine-tuning effect). Therefore, the cyclical environmental patterns, such as temperature and water inflow, end up being factored into the cyclical part of the model whilst only the residual effect is assessed by the properly named variable (e.g. temperature). Moreover, it is interesting to notice how little of the environmental variables automatically collected by the BWWTP (variable blocks “Inflow”, “V1” and “V2”) are significant to the model compared to the ones collected manually (**Figure 2b**). This may be explained by heterogeneous spatial effects, in which the surface of the tank is a patchwork of neighbouring habitats with discrepancies in parameter values, due to the viscosity of the foam. A similar microenvironment has been observed for flocks in BWWTP where nitrification was shown to happen in the outer 125 μm of the aggregates^41^.

The large importance of a “ground state” in BWWTP is linked to the need for robustness of a system that is operated primarily for public health purposes that should be hardly perturbed during parameter-controlled operations. Furthermore, it has been shown in an activated sludge population, sampled monthly over nine years, that only one out of five microbiome clusters clearly oscillated with the seasons and reached a peak abundance of 22.3% in the community^14^.

### The temporal domino of ecological events in LAO

Even if the signals S1-17 are linearly independent from one another, we hypothesised that there might be some links through time among them. These links might coalesce the system into cliques of temporally concatenated ecological events which follow each other in an ordered sequence of events (like a domino effect). We therefore used the Granger causality test, which assesses the transfer of information across time between two series of observations, to generate a causal network of S1-17 (p value < 0.05) with a maximum of ten weeks lag. The power of this representation is the amalgamation from the temporal signals, the loadings contributing to them (**Supplementary Figures 5 and 6**) and the generative model provided by ARIMA (**Figure 2b**) to generate ecological hypotheses to be further tested. Incidentally, all signals but S16, demonstrated a temporal relationship with at least another signal, resulting in a single network of causality. We decided to focus on two particular cliques of nodes in the network (**Figure 2c**) to explore the ecological domino effect: C1 (including S1, S10 andS17) and C2 (S9, S4, S7, S8).

The first clique, C1, is composed of the two “crash” signals S1 and S10, which predict each other. Indeed the peak/valley part of the signals, spanning autumn, has a similar shape but opposite sign, whilst the first part of the signals diverge, with S10 showing a sinusoidal shoulder in the beginning. Both signals are strongly dependent on their previous state in time and have clear seasonal components (**Figure 2b**). While S1 is positively influenced by four variables including the oxygen concentration as the sole environmental parameter, S10 is negatively impacted by a range of variables at the sampling site (pH, NH_4_, temperature, Dry_matter and conductivity). Podoviridae and Mimiviridae, the two virus families identified in the system, are contributing positively and negatively, respectively, to S1 in the MG (**Supplementary Figure 6**). Therefore, we infer two opposite viral mechanics involved in the fast valley to peak switch in autumn, which corresponds also to a major transient shift in community structure and substrate availability^38^. Mimiviridae target amoebas, which are known to predate on bacteria, indicating a possible multi-step, inter-kingdom curbing process. In the case of the Podoviridae, it targets Proteobacteria and Firmicutes, which are highly abundant in the LAO (**Figure 1b**). The other crash signal, S10, is characterised by the inverted reaction of the two most abundant bacterial families in the system: Microthrixaceae and Moraxellaceae (belonging to the Proteobacteria phylum). The family Moraxellaceae contributes positively to the S1 in the MG, suggesting a takeover of the community, whilst the gene expression in members of the Microthrixaceae family is repressed (negatively impact on S1, positive on S10) as shown in **Supplementary Figure 6**. It seems plausible that the rise in Podoviridae would be linked to the rise of its putative host (Moraxellaceae), to the expenses of family Microthrixaceae. However, the decrease in Mimiviridae could have triggered an increase in amoebas, resulting in greater predation on the most abundant bacterial family. These events may subsequently drive S17, a signal solely explained by a cyclic ARIMA component (**Figure 2b**), suggesting that the temporal behaviours in the systems cannot always be explained by long-term seasonal and environmental factors, but likely by the ecological interactions of the microbes involved. More specifically, S17 sees the rise in abundance or gene expression of three bacterial families: the fermenting Propionibacteriaceae, the polyphosphate accumulating Intrasporangiaceae and the autotroph Gallionellaceae. These families point to the reaction of the foam community to the observed shift in autumn. Correspondingly, S17 represents the emergence of lipid-independent metabolic strategies.

The second clique, C2, includes S9, S4 and S7 leading to S8. Both S4 and S8 represent oscillatory “perturbations” (Figure 2b, **Supplementary Figure 5d**). Whilst S4 is increasing in amplitude, S8 is decreasing. Interestingly, out of the four only S8 has an autoregressive component and S7 is missing any seasonal signal (**Figure 2b**). The nitrogen-associated S9 has a simple dependency on NH_4_ (**Figure 2b**) and indeed influences positively the family Nitrosomonadaceae (**Supplementary Figure 6**). S7 is weakly influenced by seasonality and has a relatively strong intercept (**Figure 2b**) but is affected by both pH and NH_4_. The bacterial taxonomic contributions to S7 show a mixed response of the transcriptome whereby the only positive MG association is with the viral family Mimiviridae. It is possible that S7 encodes fluctuations in the parameters and the immediate response of the microbiome (through RNA), without a defined overarching pattern. The pair S4 and S8 are however more intriguing, because of the counterintuitive idea that an escalating perturbation could contribute to the resolution of another perturbation. S4 is explained solely by seasonal components, whilst S8 also includes pH effects from both the sampling site and the inflow, even if with opposite effects (**Figure 2b**). The signal S4 is negatively associated with gene expression and protein levels, however it is positively impacted by the level of the putative predator Nannocystaceae^42^. The functional associations of S8 include a negative one for porphyrin and chlorophyll and positive ones for glycerophospholipids and simple sugars, hinting at a switch between autotrophic and medium-dependent metabolisms in the foam community. This seems to suggest that an interplay between the predation by the family Nannocystaceae, supported by parameter fluctuations in pH and NH_4_ might lead to further general instability in the RNA expression of the microbiome. Even more curious is how the exacerbation of the amplitude of S4 might drive the stabilisation of S8, according to the idea that higher predation levels have been linked to the stability of ecosystems^43^.

### Fatty Acid and Triacylglycerol accumulation are mostly time-independent

For a LAO community, the biosynthesis of Triacylglycerol (TG) and Fatty Acids (FA) are crucial steps^44^ involving multiple enzyme classes and with several entry points (**Figure 3a**). The abundant and expressed classes cover the circuit going from Acyl-Phosphate (Acyl-P) to fatty acid (FA) as shown in **Figure 3a**, however none of the enzymes’ quantities are in the top/bottom 5% of the loadings for the time-dependent EGs. It looks, in general, that the accumulation of TG and FA is time-independent. This is consistent with the observation that functions are mostly conserved in a BWWTP^14^. Interestingly K22848 is mostly encoded and expressed by the family Moraxella which is one of the two dominant families in the system **(Figure 3b**). Together with Moraxella, plasmid-encoded enzymes are also present, which was precedently unknown to our knowledge^45^, and indicates that the ability to convert DAG to TG can likely be shared between bacteria and across different taxonomic families.

**Figure 3.**
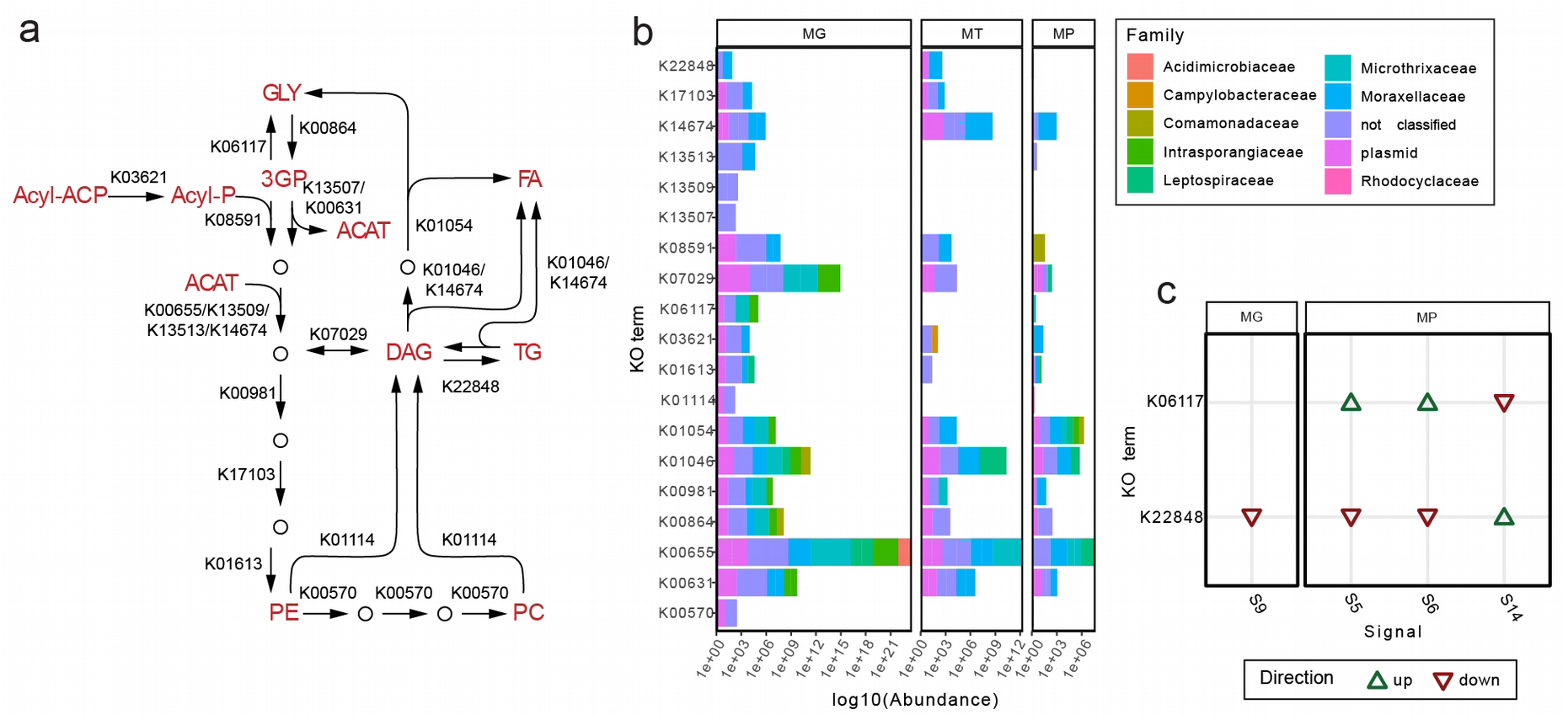
Triacylglycerol accumulation as a key metabolic community-wide trait. **a.** Enzymatic reactions (with high abundance in at least one of the omics from Shif-LAO) leading to triacylglycerol accumulation in the community. GLY: Glycerol, Acyl-ACP: Acyl Carrier Protein, Acyl-P: Acyl phosphate, 3GP: 3-glycerol phosphate, ACAT: Acetyl-CoA, FA: Fatty Acid, DAG: Diacylglycerol, TG: Triacylglycerol, PE: phosphatidylethanolamine, PC: phosphatidylcholine. The enzyme class with KO number K22848 is responsible for the conversion of DAG in TG and, ultimately, the accumulation of TG. **b.** Gene and gene product abundances for the various enzymatic groups involved in the accumulation of TG varies in amount and taxonomic origin. **c.** The gene abundance of K22484 is influenced by S9, indicating a, perhaps indirect effect on NH_4_ levels.

### Forecasting of future time points

From the analysis of the signals identified in the training datasets it is already possible to identify five signal groups: i) alternative basal states, e.g. two alternative stable states of abundance/expression, (S5, S14); ii) perturbation, i.e. standing wave with varying amplitude and frequency, (S4, S8, S15); iii) cyclical, i.e. standing wave with constant amplitude and frequency, (S6, S11, S12); iv) “crashes”, i.e. quick shift in the state and reversion to basal state, (S1, S10, S16); v) mixed, i.e. the other factors(**Supplementary Figure 6**, panels **c-f)**. Alternative stable states, perturbations and crashes (groups i, ii and iv) are hard to model without observing multiple times the shift and the perturbation events, respectively. Additionally, these scenarios may include permanent shifts into a new community equilibrium or transitory signals in the community that will be eventually resolved (e.g. a viral infection). To forecast such events and based on results, systematic information on microbial interactions would be required which is beyond the scope of this study. The used modelling also enables us to forecast cyclical events (group iii).

The 17 signals were used to train three models (with various parameters) from the package fable^37^ and the best performing model on the training set was selected for each of them (see **methods**). In detail, ARIMA, prophet and neural network models (with up to four Fourier terms for ARIMA and prophet) were trained for the S1-17 using the environmental variables as external regressors. The 51 weeks spanning 2011-2012 data were used as a training set and the model with the smallest Root Mean Square Error (RMSE) was selected for forecasting. A total of 21 new samples were collected in the month of June of the following 5 years to validate the model for the MG and MT data. To assess the accuracy of the forecasting, we computed the residues of the model and checked if they were consistent with a white noise distribution. Therefore, we showed in 16 out of 17 cases that the modelling was sufficient to reproduce the training data (**Figure 4**). The cases in which the modelling was fully successful were six: S1, S2, S4, S5, S10 and S16. However, it is worth noting that the training set for S10 was not fully captured by the model, and therefore we excluded it from further considerations. The five correctly forecasted signals account for 22.5% of EV and 24.7% of the EV by the complete S1-17 model. However, the most common outcome of the validation was a good fit to the training set and an insufficient one in the testing (9 out of 17 cases), including signals from all the groups. This could be caused by two phenomena: overfitting of the model to the training set or its insufficient size. Of particular interest is S8, whose signal in the training set remains stable for several months including the end of the training set, probably indicating that the perturbation is over. S4 is strictly tied with S8 (**Figure 2b**), however S4 was modelled and predicted correctly, suggesting that a new cycle is being established rather than a perturbation setting in. It is difficult to put these results in perspective due to the lack of similar studies covering a similar period and sampling frequency. However, the study by Wang et al.^14^, which sampled the same BWWTP monthly for nine years, showed that while five microbial clusters formed the main community, only one of them presented a clear yearly oscillating pattern. The same cluster was present in the BWWTP even after a bleaching event, therefore it is reasonable to assume that a fraction of the LAO community had a similar cluster and that the signal(s) underlying it continued in the following years.

**Figure 4.**
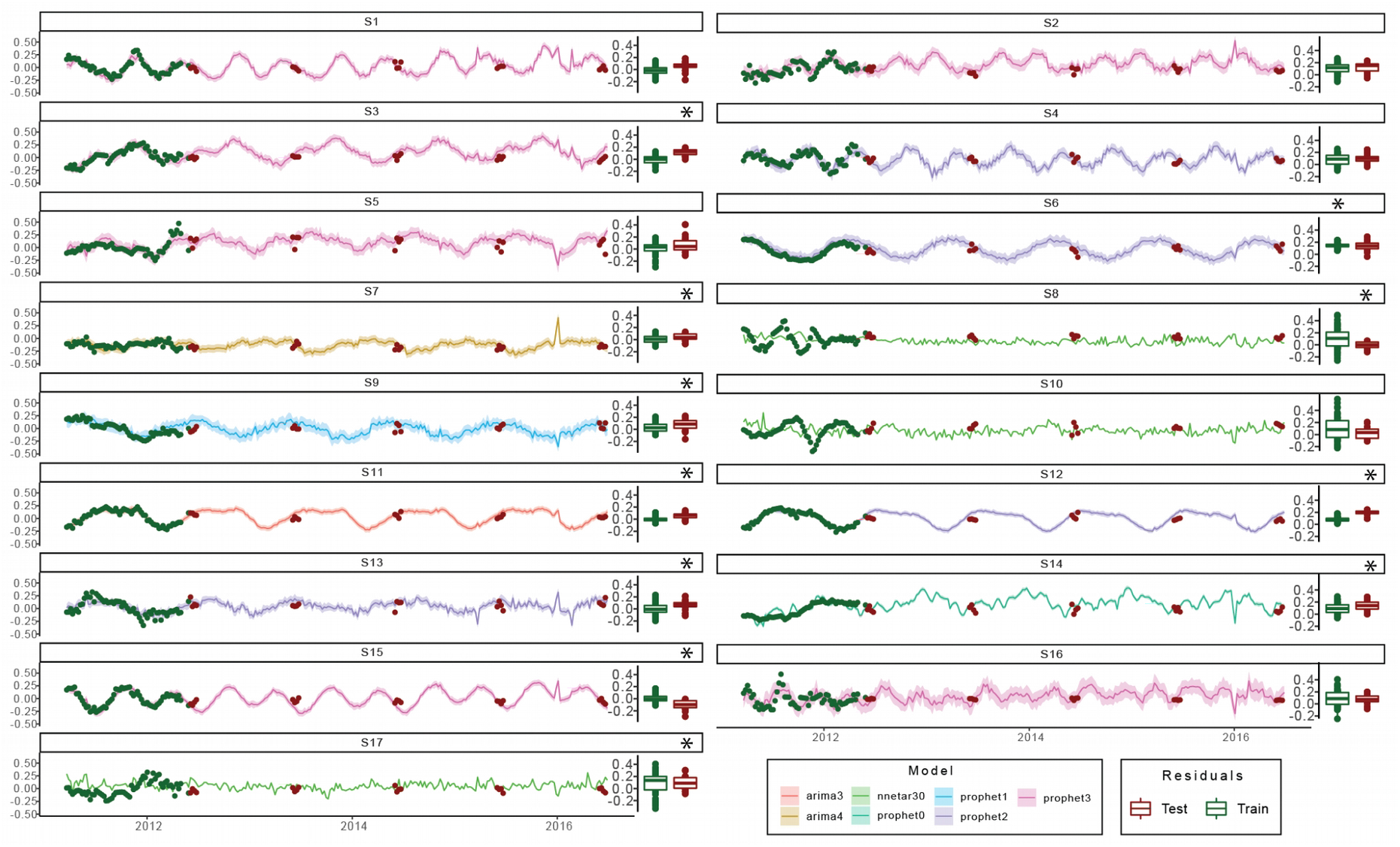
Forecasting of the signals. The 17 signals are predicted for the years 2011-2016 and compared with the data from June for those years. The green and red dots represent the training and test data respectively; the solid line depicts the median of the prediction whilst the shaded area represents the 95% confidence interval. The green and red boxplot on the right of every box depict the distributions of the model residuals from the training and the test sets, respectively. Corresponding scales are provided on the right y-axis. The residue displacement from the null distribution was assessed by a Wilcoxon two-sided test. The star on top of the boxplot indicates a statistical difference (BY corrected p value < 0.01) between the mean of the residual distribution and 0, indicating an incorrect/incomplete modelling.

Unexpectedly, the correct forecasting of S1, which looked like a crash (**Supplementary Figure 5f**) and was linked (among other things) to viral increase/decrease, suggests that it is indeed a cycle. We speculate that a recurrent triangular interaction between viruses, amoebas and bacteria might be repeated over time and lead to S1. Unfortunately, an analogous trend seen for signal S10 was not equally well represented. Similar to S1, S16 also exhibited a behaviour expected from a system crash. However, the forecasting and the testing hinted at a cyclical occurrence, hence what appeared like a crash is predicted to be constitutive and repeated behaviour. Another similarity with S1 is that viral families impacted S16, i.e. Mimiviriade (positively and negatively in the MG) and Podoviridae (positively in the MP). Signal S5 showed a sharp upward movement in relation to the general trend, before starting to dip toward the end of the time-series. Well known bacteria involved in bulking such as Moraxellaceae and Gordoniaceae have loadings contributing toward S5, hinting to a quick jolt in thickening of the foam in Summer and an overall cyclical effect that can be forecasted over time.

### Forecasting gene abundance and expression

Following the forecasting of the signals, we decided to try to reconstruct the samples of the following years. The signals harbour (almost) all the temporal information but they need to be weighted again to re-write the LAO samples according to them. Therefore, we re-wrote the abundance and gene expression of the microbial families as well as reactions (KO term groups) and pathways in the terms of the S1-17 using a linear model. We then reconstructed those matrices for the test sets using the S1-17 forecastings and the weights of the models as well as the intercepts and compared the reconstructed values with the original ones for each sample (**Supplementary Figure 10**). The comparisons show a range of results including samples that were predicted correctly (data points arranged in a narrow diagonal line), samples with poor predictions (unordered distribution of the data points) and samples with an unexpected inverse relationship with the prediction (descending diagonal line). When taking into account the explanatory variables in the ARIMA modelling we already hypothesised a micro-environmental effect at play in the foam, making it a composition of areas with (slightly) different environmental values. We now extend that hypothesis to the sampling unit itself (the foam “islet”, see **Methods**), which might have individual genetic potential and gene expression characteristics imputable to the process of foam formation, permanence and stability. We therefore assume that the islet variability, compounded with the temporal evolution of the system, has ultimately an impact on the sample. Intuitively, if the foam islets were composed of the same genetic makeup but subject to (even small) different environmental conditions, one would expect gene abundances to be relatively stable yet gene expression might change. Instead, observing the coherent response between MG and MT to the reconstructed samples from **Supplementary Figure 10,** it is apparent that the genetic makeup of the islets changes from week to week and gene expression changes accordingly to this alteration. We assume that our modelling creates a “smoother” representation of the data, necessarily averaging the observed sample to sample variability. This can be imputable to the SVD step of the modelling, which isolates “high level” patterns that harbour lower noise than any individual ORF- or descriptor-based summarisation of the data. Moreover, the scale of the values is often larger in the reconstructed samples than in the test ones (**Supplementary Figure 10**).

To counter the islet variability, we considered the average of the measured and predicted values over the month of June for each year and computed the coefficient of variation R^2^ for each of them (**Figure 2**). The R^2^ is strikingly high (≥0.97) in all the six matrices for the first two years following the training set but the predictability starts decreasing from the third year after the last sample. This implies that in our system (Shif-LAO) the observation through multi-omics data and the environmental parameters for 14 months is sufficient to build a reliable predictive model. Moreover, with this model and the monitoring of the environmental parameters, it is possible to correctly chart the community structure and function at any given point within the two years following the sampling.

## Conclusions

We present the temporal reconstruction of the surface microbial community of a BWWTP over a year and a half of weekly sampling. The gene abundance and expression show 17 distinct and linearly independent signals (S1-17) across time (**Figure 2a**), many of which were explained by the physicochemical parameters and the mathematical components describing self-dependence and seasonality (**Figure 2c**). The signals were tied in a “temporal domino” (**Figure 2b**), from which we selected two cliques to successfully describe: the “autumn crash” (C1) and an oscillatory perturbation (C2). The models built on the S1-17 signals and paired with the environmental parameters, were subsequently used to forecast the next five years of the LAO community. We demonstrate that five of the forecasted signals (S1, S2, S4, S5 and S16) are indeed validated by the future samples (**Figure 4**) and cover some interesting aspects of the BWWTP surface community like Nitrogen metabolism (S4) and viral interplay (S1 and S16), as well as well-known foam-related dynamics (S5). Importantly, when rebuilding the gene abundance and expression data at the levels of taxonomic families, reactions and pathways and extrapolating to the future samples (June 2012-2016) the results over the averaged month of June were near perfect for the first two years after sampling (R^2^≥0.97). However, a clear fading was apparent starting from the third year (**Figure 5**).

**Figure 5.**
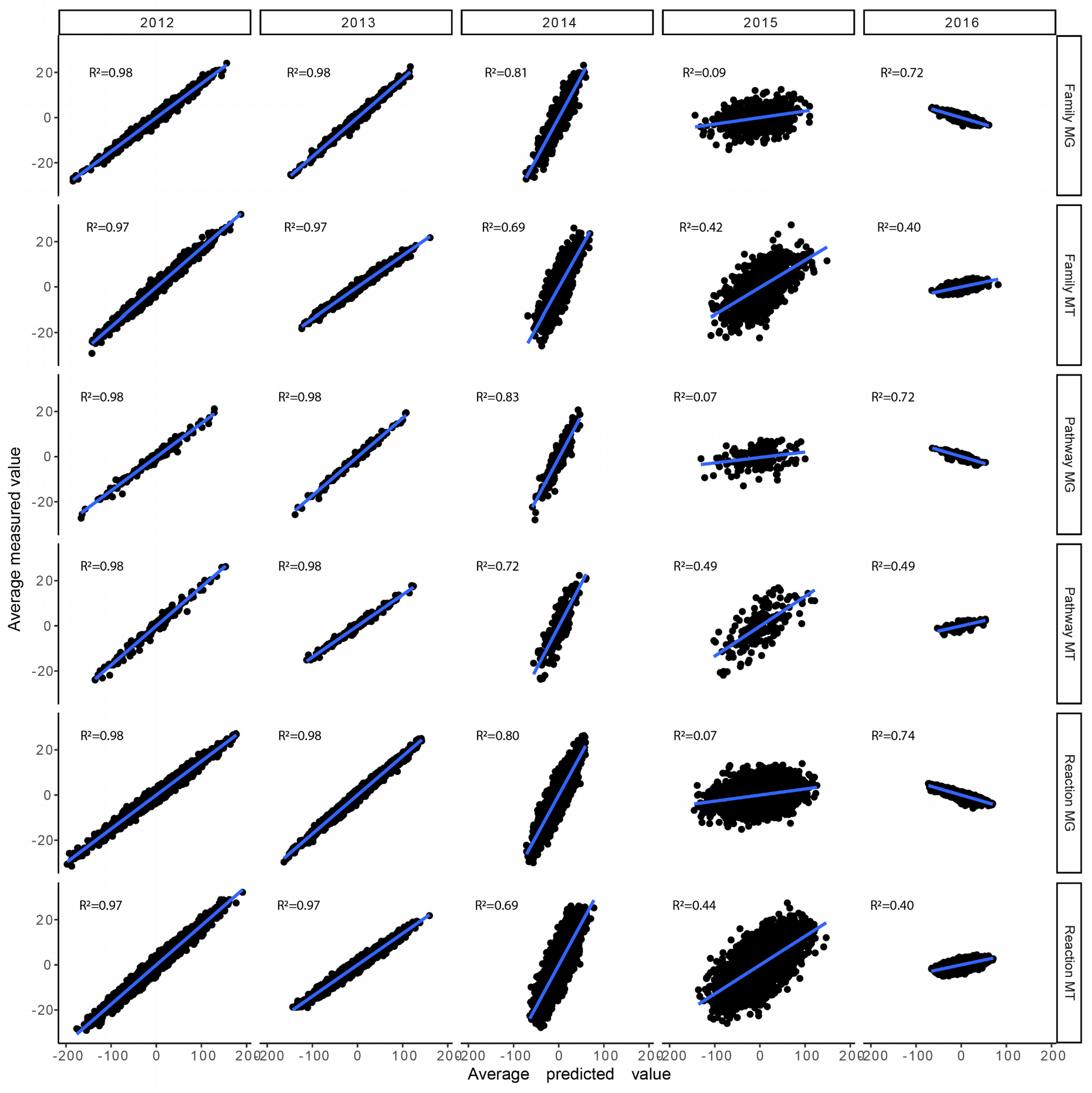
Reconstruction of the June months 2012-2016. The test samples were reconstructed using the 17 signals and their weights estimated through linear regressions on the training set. The reconstructed matrices are based on MG and MT data summarising taxonomic families, reactions and pathways. The coefficient of determination R_2_ is reported for each panel with a higher coefficient demonstrating a more accurate prediction.

Overall, the present approach covers the vast majority of time-dependent information in the system. It furthermore enables us to describe a complex community with its behaviour in a number of temporal patterns which is easy for a human to interpret (in our case 17 signals) and link these to their underlying generative processes, as well as the environmental parameters, taxa and functions supported by them. Furthermore, the method allows to reliably forecast these fundamental signals that represent a seasonality and temporal span (more than one year, hence more than one expected full cycle of the system), indicating that the time- and environment-dependent components can explain the community during regular WWTP operations. We hope that further work, especially the sampling the BWWTP at higher time frequencies (e.g. hours) and/or for longer periods (multi-annual training sets), could be integrated for a more detailed systemic description and increased ability to forecast in order to cover those phenomena that our work falls short of. Finally, we infer that there are environmental drivers in the macroscopic composition of the LAO community behaviour and that we are able to correctly reconstruct the samples from 2 years in the future but that the foam presents a high islet variation which is beyond the predictability of this method.

## Data and code availability

The generated MG and MT reads (FASTQ) files, as well as the previously produced data, are available as NCBI BioProject PRJNA230567. The MP data from the PRIDE repository with accession number PXD013655^38^.

The meta-omics pipeline IMP v3.0^46^ is maintained and developed at the GitLab page: https://git-r3lab.uni.lu/IMP/imp3. The code used in the analysis is available at https://github.com/fdelogu/microforecast, whilst the data required to start the analysis is available on Zeonodo with the doi 10.5281/zenodo.7225349.

## Materials and Methods

### Sampling and preprocessing

Floating LAO biomass was sampled from the air–water interface of the anoxic activated sludge tank at the Schifflange wastewater treatment plant (Esch-sur-Alzette, Luxembourg; 49130048.2900N; 61104.5300E) in the form of a single islet (examples illustrated in **Figure 2** from Roume et al.^39^). The sampling frequency - weekly- was chosen as it is the generation time of the activated sludge in the BWWTP (the average time it remains in the system) and the average doubling time of the dominant Microthrix population^47^. For each sampling date, indicated as dates in the format YYYY-MM-DD, one entire ‘islet’ was sampled using a levy cane of 500 ml. Samples were quickly homogenised and collected in 50 ml sterile Falcon tubes and then immediately flash-frozen by immersion in liquid nitrogen and stored at −80°C to guarantee optimal sample integrity and quality.

For the 51 time points of the training set (2011-03-21 to 2012-05-03), were treated in 2012 as previously described^38^: 200mg were subsampled from the collected islet using a sterile metal spatula while at all times guaranteeing that the samples remained in the frozen state and used for subsequent biomolecular extraction according to previously published procedure (using the Qiagen AllPrep DNA/RNA/Protein Mini kit-based method on “LAO-enriched mixed microbial community”^35^).

Additional concomitant biomolecular extractions were applied to a total of 21 samples collected during the month of June from 2012 to 2016, and extracted in a separate experiment in 2018. The sample pre-processing protocol has been carried out on a customised robotic system owned by the lab (Beckman-Coulter_Platform Biomek 4000 NXP Span8 Gripper) following the same protocol as for the training set sample extraction described above with few differences. The biomolecular extraction was then performed using the commercial AllPrep DNA/RNA/Protein Mini Kit (Qiagen-80004), conducted on a customised robotic system owned by the lab (Tecan-LU_UNILU_EWS_EXTRACTION_EU-0908-Freedom EVO 200). A RNase treatment followed by DNA precipitation was carried out on the DNA and the RNA was purified by using the commercial kits Zymo RNA Clean&Concentrator-5 (Ref: R1013). RNA quality was assessed as in the previous study for the same environment^39^.

### High-throughput meta-omics

400 ng of DNA was sheared using NGS Bioruptor (Diogenode, UCD300) with 30s ON and 30s OFF for 10 cycles. DNA libraries were prepared using TruSeq Nano DNA kit (Illumina, FC-121-4002) using standard protocol with 8 PCR cycles. The libraries were prepared for 350bp average insert size. 1μg of RNA was rRNA depleted using the RiboZero kit (Illumina, MRZB12424). rRNA depleted samples were further processed and prepared using TruSeq Stranded mRNA library preparation kit (Illumina, RS-122-2101). The fragmentation time was reduced to 3min. The samples were amplified for 8 PCR cycles. The prepared libraries were quantified using Qubit (ThermoFischer) and quality checked using Bianaoyzer 2100 (Agilent). Sequencing was performed on NexSeq500 instrument using 2×150bp read length at LCSB sequencing platform (RRID SCR_021931).

### Collection of environmental variables

The environmental variables were collected on site by the researcher(s) whilst they were performing the sampling, which include: dry matter, phosphate, nitrate, ammonium, oxygen, conductivity, pH, temperature and oxygen (**Supplementary Table 4**) following the previously established protocol^38^. The other variables were retrieved from the automated data collection routine of the Schifflange BWWTP, which measures online these values and aggregate them as 2h average starting at 1:00 am. Those recordings include the same variables for different parts of the plant (inflow, both vats, outflow) with the addition of other measurements such as the in/out flow volume. For simplicity, we used exclusively the variable pertaining to the inflow, both vats and outflow in this study (**Supplementary Table 5**). The Schifflange plant is depicted in https://sivec.lu/installation/station-depuration/ with the various components named in german. The variables were screened for collinearity (**Supplementary Figure 2**) using a Pearson Correlation Coefficient threshold of 0.7. For each cluster of correlated variables a single one was selected, resulting in 15 variables used from the 59 initial ones. The variables Oxygen_manual, Dry_matter, NH4.N, Vat1_NH4.N, Vat2_NH4.N were transformed using the square root function.

### Co-assembly of metagenomics and metatranscriptomics reads

All the samples from the training and the test datasets followed the same bioinformatic pipeline. Sample-wise preprocessing of the MG and MT data was performed using IMP v3.0^46^ (https://git-r3lab.uni.lu/IMP/imp3) with custom parameters, i.e. i) Illumina Truseq2 adapters were trimmed, ii) the step involving the filtering of reads of human origin step was omitted for the preprocessing. The reads were corrected using BayesHammer^48^ per sample, per omic. The resulting MG and MT reads were assembled with metaSPAdes v3.13.1^49^ and rnaSPAdes v3.13.1^50^ respectively. The MG and MT reads of each sample were re-assembled together using the contigs and “highly filtered” transcripts from the first assemblies as trusted contigs.

### Contig sorting into biological subsets

The contigs longer than 1000 nt from each sample were retained and were sorted into four subsets: eukaryotes, plasmids, viruses and chromosomal prokaryotes. First the contigs were screened for eukaryotes using EukRep^51^; the resulting non-eukaryotic contigs were searched for plasmidial sequences with Plasflow^52^ and cbar^53^ as well as for viral sequences using virsorter (category 1 and 2)^54^ and deepvirfinder^55^. A contig was considered viral or plasmidial if both tools agreed in the prediction, all the leftover sequences were considered chromosomal prokaryotic. Later some contigs of the latter group were moved to the eukaryotic (see the **Taxonomic annotation** section).

### Binning and clustering

The chromosomal prokaryotic subsets of each sample were binned using IMP v3.0^46^ with MaxBin^56^, MetaBAT^57^, binny^58^ plus a refinement step with dastool. The resulting bins dereplicated along the entire time series with dRep^59^ to create representative metagenome assembled genomes (rMAGs). Similarly, the eukaryotic subsets were binned with MetaBat^60^ and dereplicated using dRep^59^ without genome quality assessment resulting in rMAGs. All the plasmidial, viral and the unbinned contigs from the eukaryotic and chromosomal prokaryotic subsets were clustered using CD-HIT^61^ on each of those subsets. We refer to the subset of the clustered unbinned contigs as representative contigs (rContigs). The collection of the rMAGs and the rContigs constitute the representative database (rDB) of the system.

### Taxonomic and functional annotation

The rMAGs and the rContigs were annotated taxonomically using CAT and BAT^62^ respectively. The ORFs were predicted from the rDB using IMP v3.0^46^ and annotated using Mantis v1.02^63^ with the heuristic approach and using all the databases. Subsequently only the entries with KO terms assigned by kofam were retained for analysis.

### MG and MT quantification and filtering

The filtered MG and MT reads were aligned to the ORF reference set using bwa^64^ and sorted using samtools^65^. The resulting sorted bam files were processed using bam2hits^66^, and the output split with a maximum number of 100’000 ORFs per subset, whilst respecting the bam2hits read groups. Each subset was quantified with mmseq^66^ and mmcollapse^67^, then the quantifications per sample were the-normalized form FPKM, merged and re-normalized to FPKM. Values of gene abundance and expression inferior to 10^−7^ were considered equal to 0 and ORFs and transcripts that were not present in at least 20% of the training set were discarded from further analysis.

### MP quantification and filtering

Raw MP data were retrieved from the PRIDE repository with accession number PXD013655^38^, where the samples were processed as described in Muller et al.^47^, and we reanalyse them. Supplementary Figure 10The complete set of predicted ORFs was subsetted to obtain smaller sample-specific databases. The MG alignment files generated in the previous step were processed with featurecounts^68^ and all the ORFs with a count greater than 0 for the given sample were included in the appropriate sample. Each sample-specific database was concatenated with a cRAP database of contaminants (https://thegpm.org/cRAP; downloaded in July 2019) and the human UniProtKB Reference Proteome (UniProt Consortium, 2021), and decoys were generated by adding the reversed sequences of all protein entries to the databases for the estimation of false discovery rates. The search was performed using SearchGUI v. 3.3.20^69^ with the X!Tandem^70^, MS-GF+^71^ and Comet^72^ as search engines and the following parameters: trypsin was used as the digestion enzyme and a maximum of two missed cleavages was allowed. The tolerance levels for matching to the database were 10 ppm for MS1 and 15 ppm for MS2. Carbamidomethylation of cysteine residues and oxidation of methionines were set as fixed and variable modifications, respectively. Peptides with length between 7 and 60 amino acids, and with a charge state composed between +2 and +4 were considered for identification. The results from SearchGUI were merged using PeptideShaker-1.16.45^73^ and all identifications were filtered to achieve a protein false discovery rate (FDR) of 1%. The sample-specific peptide-spectrum matches (PSM) obtained for each analysis were then used to calculate dataset-wide protein groups using the Occam subgroup method from the Pout2Prot algorithm^74^. The dataset-wide protein group output was then submitted to Prophane^75^ with default parameters to retrieve the quantitative values using normalised spectral abundance factor (NSAF). Values of protein abundance inferior to 10^−3^ were considered equal to 0 and only proteins present in at least 20% of the training samples were retained for further analysis.

### Batch effect correction

The whole data analysis was conducted in R 3.4.4. Firstly we transformed the MG, MT and MP data using the central log ratio with the function *clr*^76^ to overcome the inherent problems of compositional data^77,78^. In order to estimate the batch effect between the train and test samples, introduced by the different experimental procedure (mainly the robotic biomolecular extraction in the test samples and the read length), we regressed every entry in the MG and MT matrices with a linear model (with the function *lm*) as:

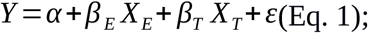

where Y is the central log ratio (clr) transformed quantification matrix, α is the intercept of the model, X_E_ and X_T_ are the environmental and technical variables (number of reads, average length of reads), respectively, β_E_ and β_T_ are the vectors of the environmental and technical coefficients, respectively and ε is the randomly distributed gaussian error N(0, σ^2^). The non-normality of β_T_ was assessed with the shapiro test^79^ (function *shapiro.test)*, sampling 10 times 5000 ORFs at random per technical variable for the MG and MT matrices respectively and computing the scores in **Supplementary Table 2** and **3**. Therefore we corrected the quantification matrices as:

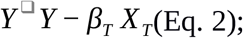

subtracting the estimated batch effect from the quantification matrices. The distributions of the β_T_ are shown in **Supplementary Figure 1**.

### Eigengenes and their analysis

The EGs for the training set (samples from 2011-03-21 to 2012-05-03) were computed as singular right eigenvectors obtained with the function *svd*. The data were normalised according to the basal expression^27^ computing the quantification matrices as:

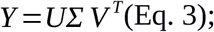

where the first element of the eigenvalues vector Σ has been replaced by 0. The EGs were recomputed from the normalised matrices and subsequently tested using the Ljung-Box test *(Box.test)*, the augmented Dickey-Fuller test (*adf.test*) and the Kwiatkowski–Phillips–Schmidt–Shin (*kpss.tests*) tests with null hypotheses “trend” and “level”. If at least two of the four tests were passed (p<0.05 for Ljung-Box and Kwiatkowski–Phillips–Schmidt–Shin tests; p> 0.05 for Dickey-Fuller test) the EG was considered time-dependent. The i^th^ EG was modelled using seasonal ARIMA modelling. Considering that the training set did not span two cycles (the hypothetical period of seasonal patterns) we added up to 4 Fourier terms to the model as a proxy for the seasonal component. We used the *arima* function from the package fable^37^ as:

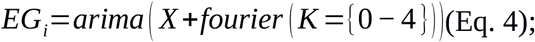

where X is the matrix of the environmental variables and the Fourier term includes 0 to 4 components. The best model of the five was selected according to the R^2^ value of the models. Finally, we assessed the significance of the explanatory variables using ANOVA (*anova*).

### Eigengenes clustering and Granger causality network

Correlations between pairs of EGs were computed with Pearson linear correlation, the output was made absolute and the minkowski distance was computed. The clusters were retrieved using the *cutreeDynamic* function (deepSplit=0, pamRespectsDendro=FALSE, minClusterSize=) from the dynamicTreeCut package^80^, resulting in 17 groups (**Supplementary Figure 5**). From each of the 17 groups a representative EG was selected according to the following criteria: i) MG or MT (because MP data do not exist beyond the training set), ii) smoothest profile (minimal median of the absolute de-trended time series). The resulting EGs are the S1-17 in **Figure 2a**.

The signals were tested two at the time with the Granger causality test (grangertest) from the lmtest package^81^ and if the p-value was inferior than 0.05 two signals were considered connected. The visualisation of the network was performed with Cytoscape^82^.

### Signal forecasting and gene abundance/expression reconstruction

For each signal we tested ARIMA with up to four Fourier components, Prophet with up to four seasonal components and a neural network with 10, 20 and 30 nodes in the hidden layer. The models were scored according to their RMSE and the top three were combined (weighted by 1-RMSE) and used as a fourteenth model, for which the RMSE was calculated too. The best of the fourteen models was selected for each signal and used to forecast the test test with the function forecast form the fable package and supplying environmental parameter readings.

All the 27 matrices used to summarise the LAO community can be rewritten using a linear combination of the 17 signals plus a basal abundance/expression (removed in the analysis). We therefore decided to rebuild June 2012-2016 matrices for the reaction, pathway and family summarisation of the gene abundance (MG) and expression (MT). We run linear regression (lm function) using the six training set matrices for the categories above as target variables and the 17 signals as explanatory variables. We then rebuild the test matrices multiplying the forecasted signals over the test set time with the newly calibrated betas and adding the intercept (basal level). The reconstructed samples were compared with the original ones on an individual basis (**Supplementary Figure 10**) and on an average one (**Figure 4**).

## Author contributions

F.D. and P.W. contributed to the planning and designing of the overall study and analyses. F.D. performed the data analyses. P.M.Q. contributed with the protein annotation software. B.J.K. performed the MP measurement. E.E.L.M. and L.A.L. collected and performed the biomolecular extractions on the samples. R.H. performed the DNA and RNA sequencing. F.D., P.M., S.W., E.E.L.M. and P.W. participated in discussions related to this work. F.D., P.M., S.W., E.E.L.M. and P.W. wrote and reviewed the manuscript. All authors read and approved the final manuscript.

## Acknowledgments

We thank the Luxembourg National Research Fund (FNR) for supporting this work through various funding instruments. Specifically, a PRIDE doctoral training unit grant (PRIDE/15/10907093), CORE grants (CORE/17/SM/11689322), a European Union ERASysAPP grant (INTER/SYSAPP/14/05), and an ATTRACT grant (A09/03) all awarded to P.W. as well as CORE Junior (C15/SR/10404839) grant to EELM. The project received financial support from the Integrated Biobank of Luxembourg with funds from the Luxembourg Ministry of Higher Education and Research. This work was also supported by the European Research Council (ERC) under the European Union’s Horizon 2020 research and innovation programme (grant agreement No. 863664). The work of P.M. was funded by the ‘Plan Technologies de la Santé du Gouvernement du Grand-Duché de Luxembourg’ through the Luxembourg Centre for Systems Biomedicine (LCSB), University of Luxembourg. S.W. was supported by the Austrian Science Fund (FWF) Elise Richter V585-B31. P.B.P is grateful for support from The Research Council of Norway (FRIPRO program: 250479) and The Novo Nordisk Foundation (Project no. 0054575). The authors acknowledge the ULHPC for providing and maintaining the computing resources. We duly thank Mr. Bissen and Mr. Di Pentima from the Syndicat Intercommunal a Vocation Ecologique (SIVEC), for access to the Schifflange wastewater treatment plant.

