## Supplementary Material for "Forecasting of a complex microbial community using meta-omics"

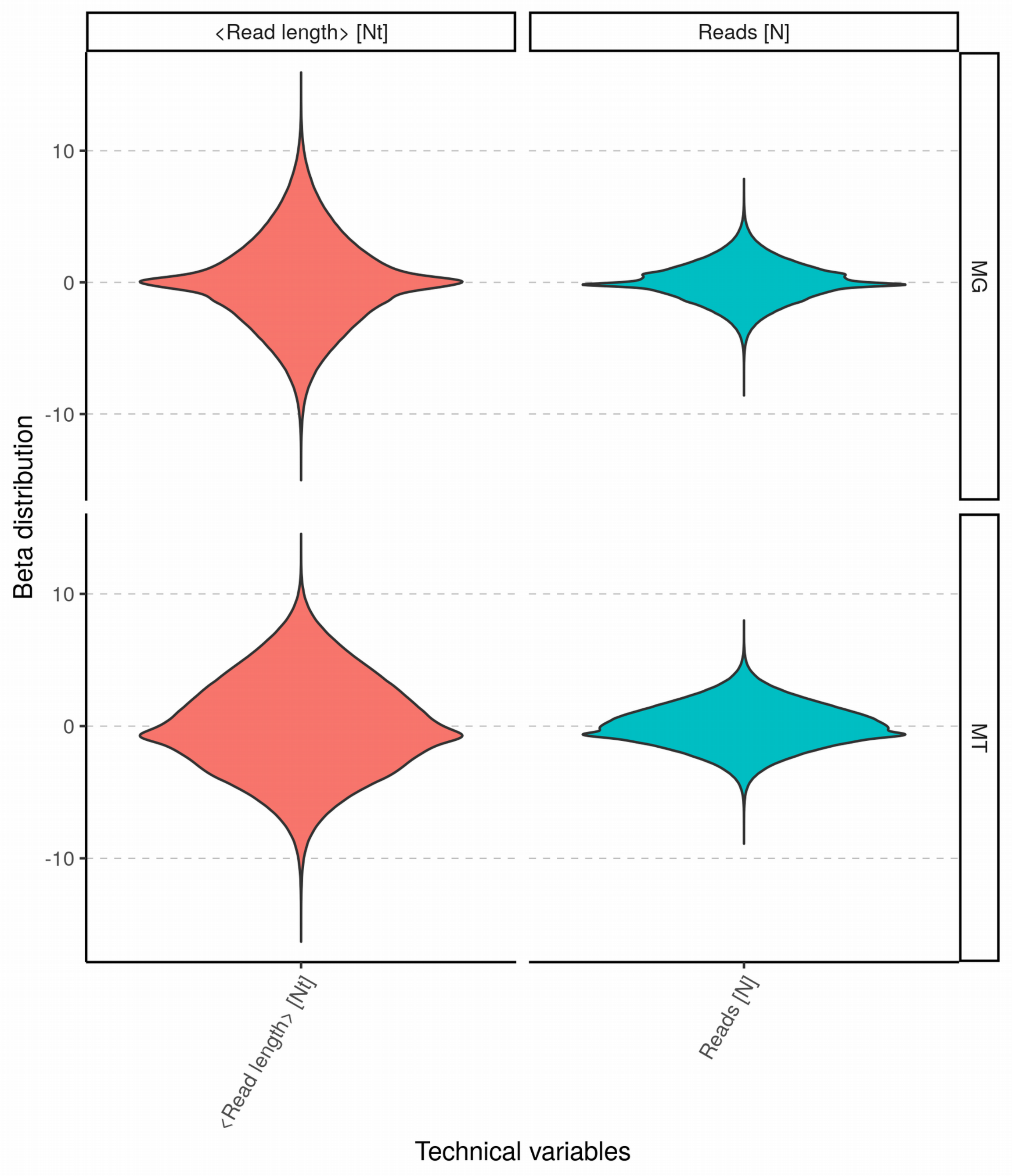


**Supplementary Figure 1**. Technical effect estimation. The data were regressed with the experimental variables (i.e. environmental parameters) and the technical ones (i.e. read length and number of reads). The plot shows the distribution of the betas resulting from the regression for the MG and MT ORF-based matrices.

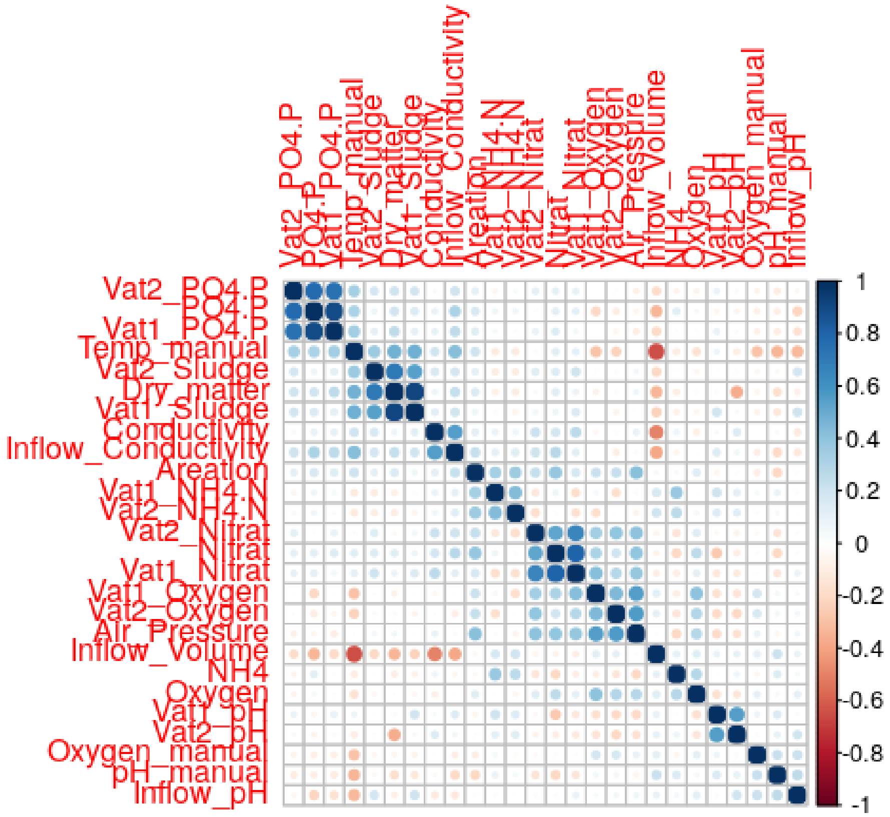


**Supplementary Figure 2**. Corr-corr plot of the correlations between the selected starting environmental variables to explain the signals. Form here the final 16 variables were selected.

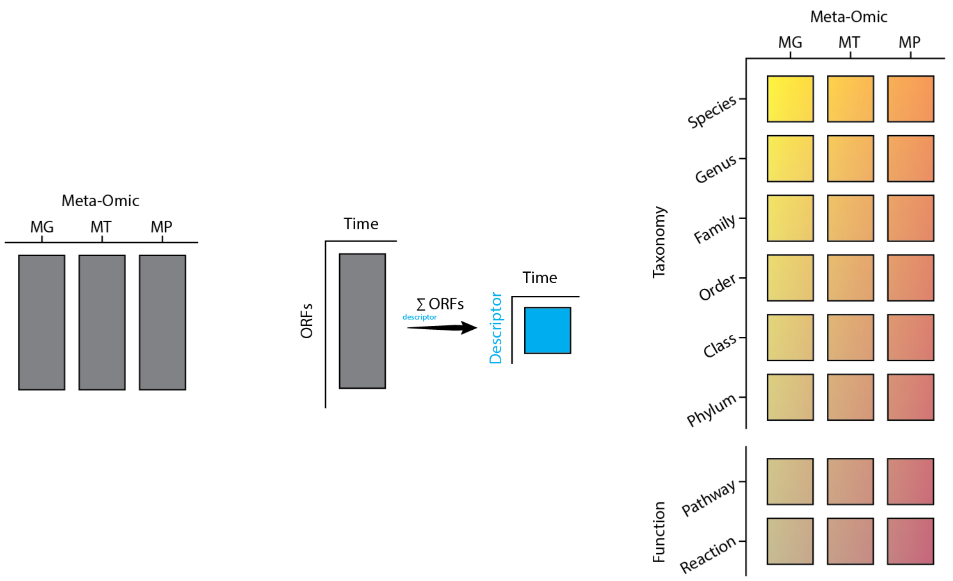


**Supplementary Figure 3**. The three ORF-based omic quantification matrices are summarised by summing up the lines with the same ORF descriptor. The final result is a collection of 24 matrices + the original three.

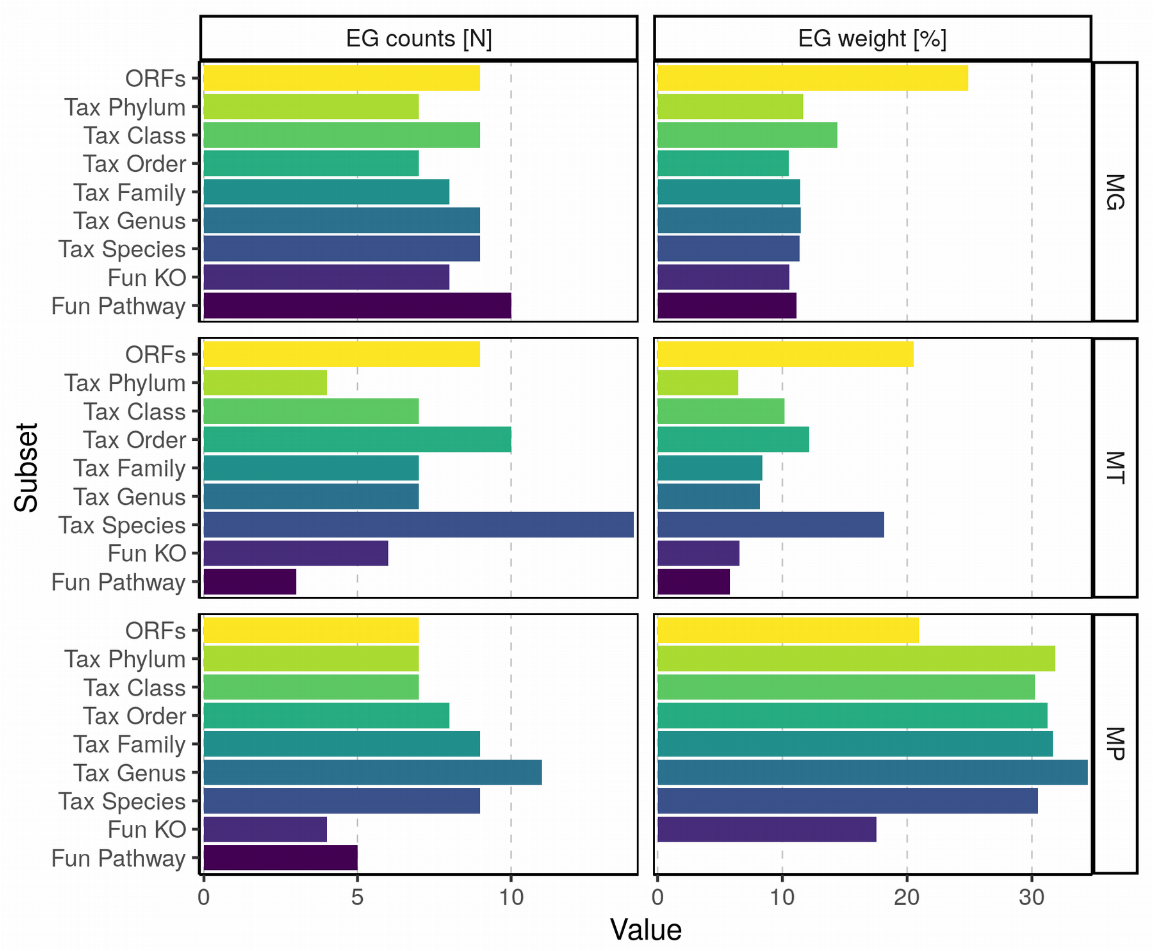


**Supplementary Figure 4**. The six panels show the number of time-dependent EGs and the EG weights (equivalent to the Explained Variance) per omic in the nine summarisation matrices. The first EG (i.e. the basal state of system) was removed and all the EGs re-scaled per matrix. In the y axis "Fun" stands for "Function" and "Tax" for "Taxonomy". The number of selected EGs changes depending on the omic and the descriptor, however some trends can be seen in the EG weight. For MG and MT the EG weight is the largest, signifying that it is, if taken alone, the most informative layer of information. Interestingly in MT the second largest, with a decent margin, is the Species level, which can be explained as a level in which most of the individual genes information is conserved (i.e. genes of the same species will be expressed together over time).

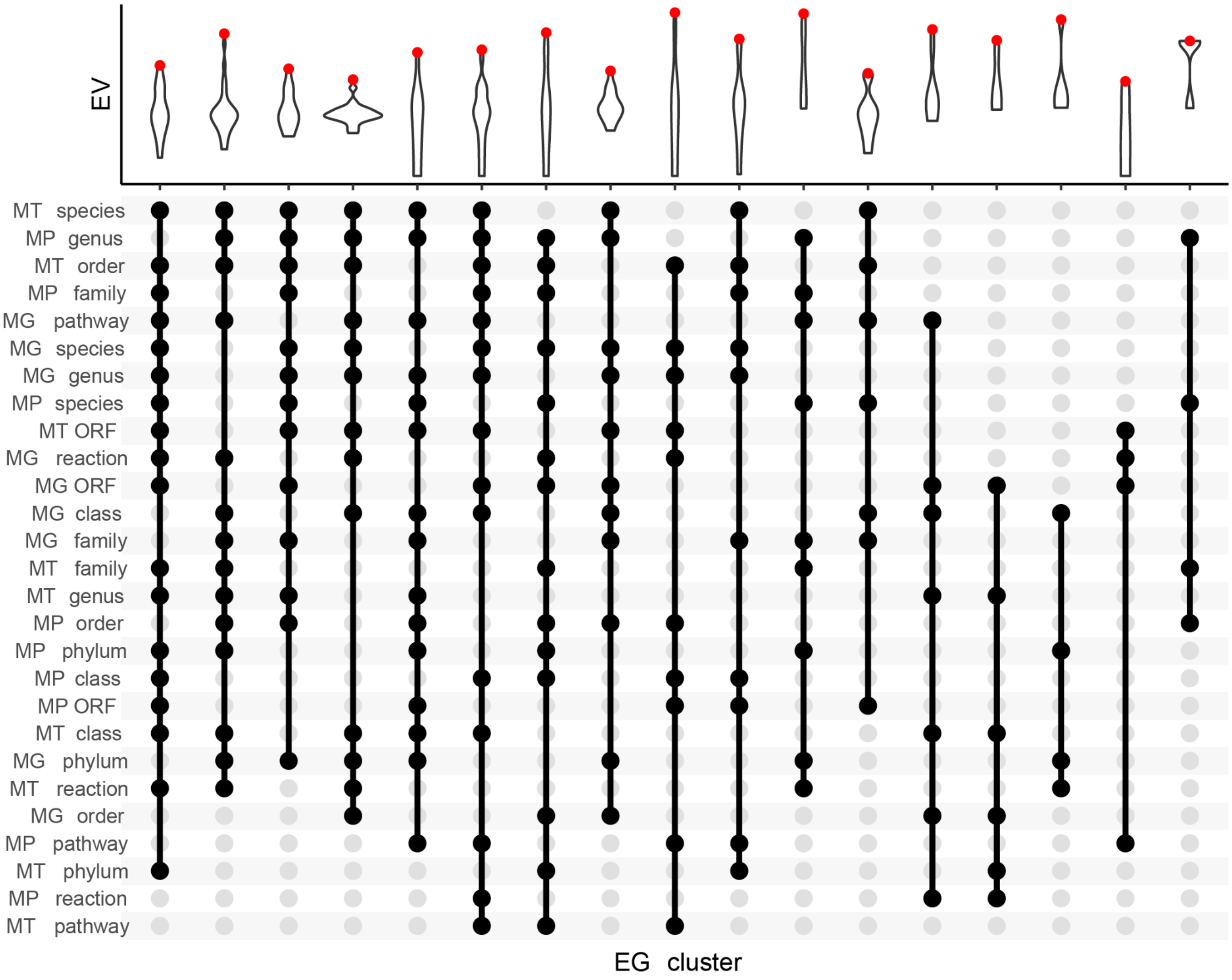


**Supplementary Figure 5**. EG clustering. The columns represent the 17 EG clusters while rows indicate the different types of summarisation matrices. In the top panel the violin plots depict the distribution of the explained variance (EV) from the EGs in the cluster. The red dot indicates the maximal EV in the distribution and the EV of the cluster. On the y-axis there are the 27 matrices.

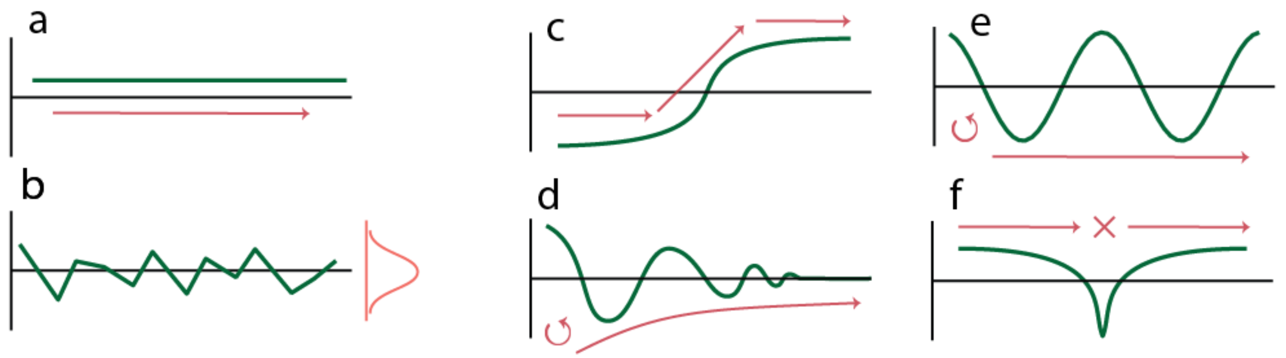


**Supplementary Figure 6**. Example of 6 patterns detectable in time series. **a.** Basal level, like the one excluded by removing the first EG in the analysis; **b.** Random noise; **c.** level change; **d.** perturbation; **e.** cycle; **f.** Crash. In real time series more patterns are usually combined (at least with noise) to create the main data behaviour over time.

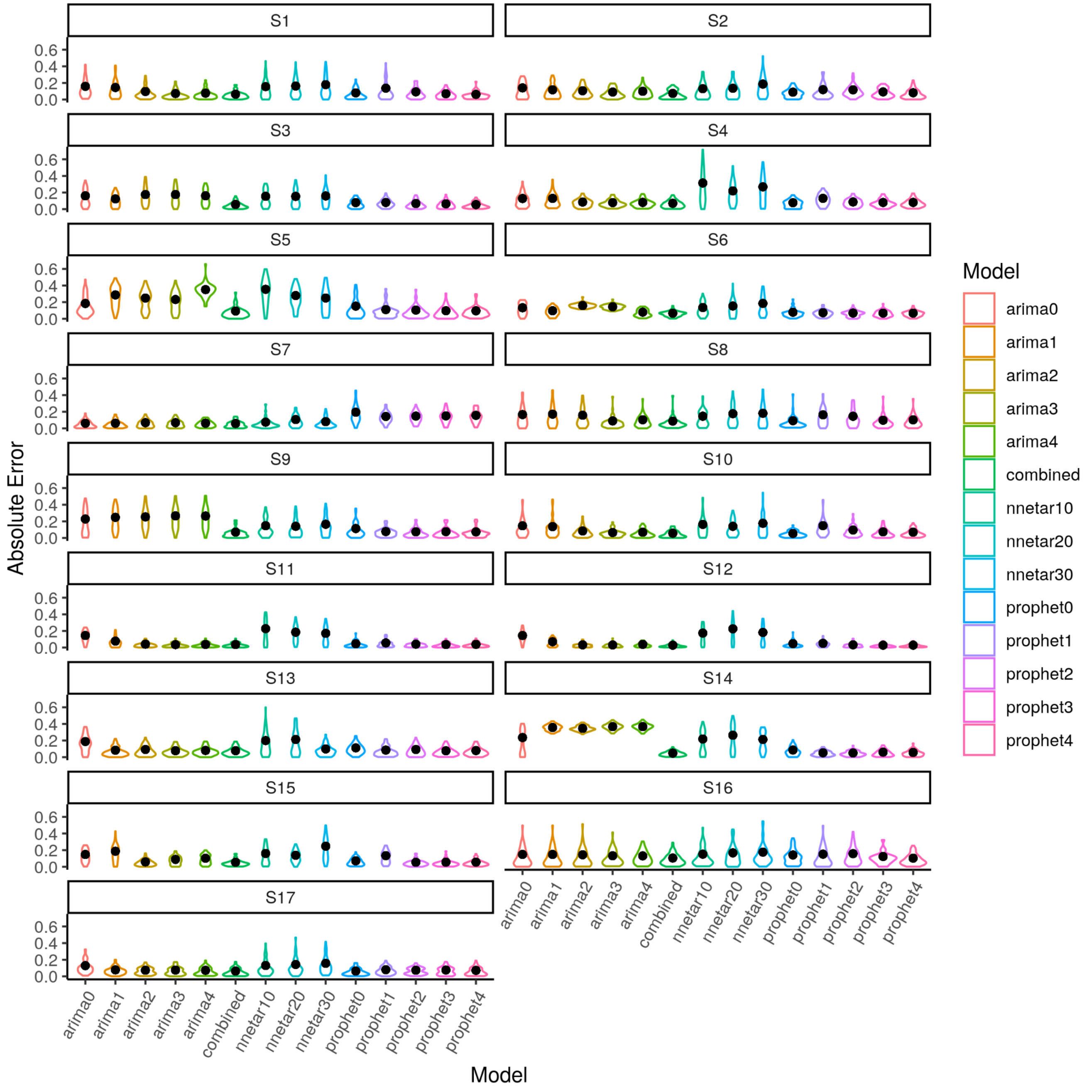


**Supplementary Figure 7**. Absolute error profiles over the training data of the eleven tested models for each of the seventeen signals S1-17; the black dot indicates the RSME. A low RMSE indicates that the predictions and the real data are close; vice-versa a high value shows distant data points. Therefore RMSE is useful when comparing multiple models. Elongated violin plots indicated a spread of values (i.e. both correctly and incorrectly predicted weeks), a "short" and "wide" distribution with an upper tail indicated a "focused" prediction overall with some outliers, whilst a simple "short" and "wide" distribution is obtained for very coherent predictions (i.e. constantly correct or incorrect). The dots represent the average RMSE.

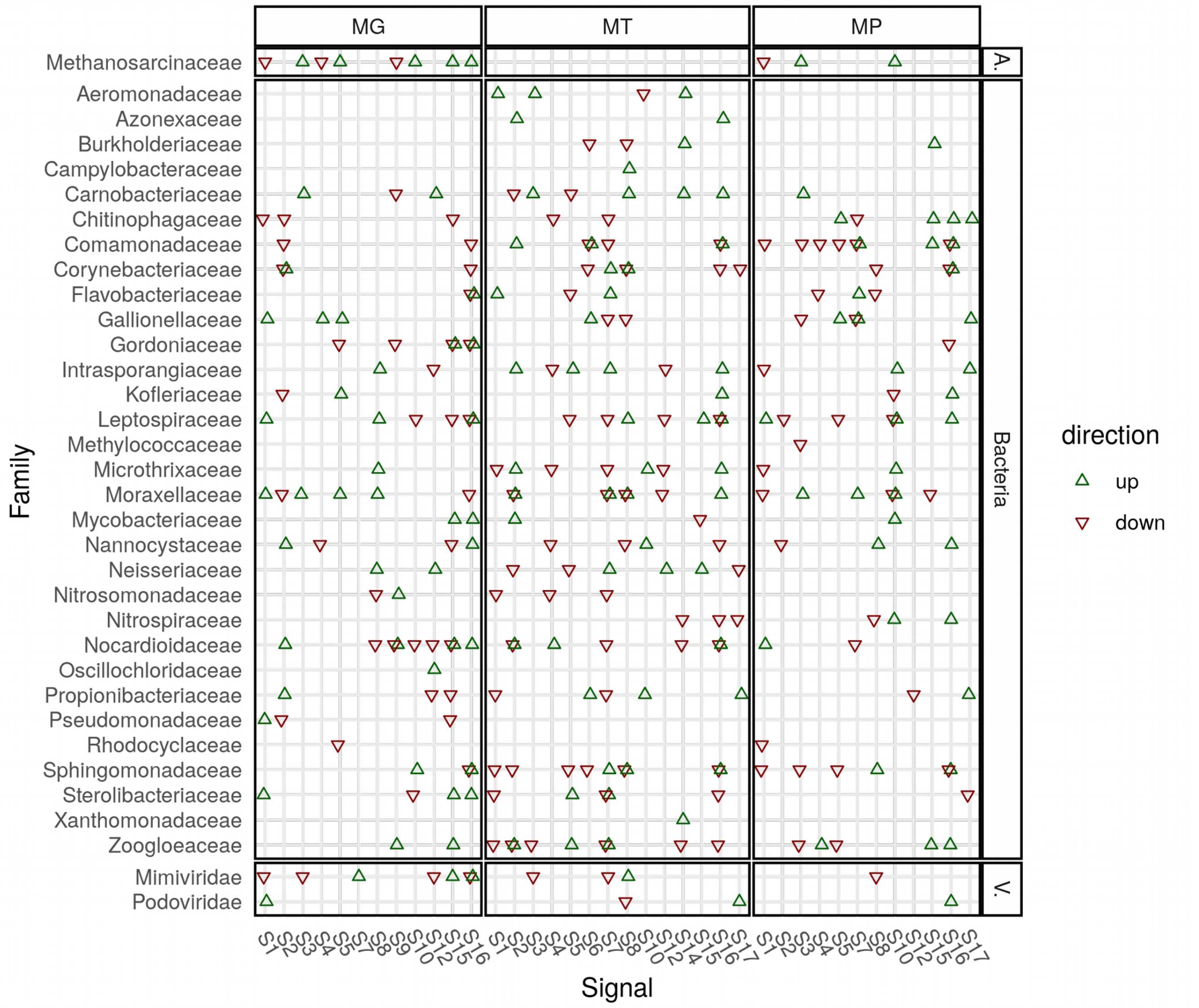


**Supplementary Figure 8**. Loadings at the family level. On the y-axis the taxonomic families intersect the signals they contribute to from the x-axis. If the loading is in the top 5% a green arrow pointing up marks the intersection. Similarly if the loading is in the bottom 5% (strongly negative) a red pointing down marks the intersection. The vertical blocks separate the three omics, whilst the horizontal blocks separate the archaea (A.), bacteria and viruses (V.). No eukaryotic families were found to be in the top/ bottom 5% of the loadings. The plot also integrates lower taxonomic labels (i.e. species and genus) and some of them might have opposite orientations, leading to families with both types of arrows.

**Supplementary Figure 9**. Loadings at the pathway level. On the y-axis the metabolic pathways intersect the signals they contribute to from the x-axis. If the loading is in the top 5% a green arrow

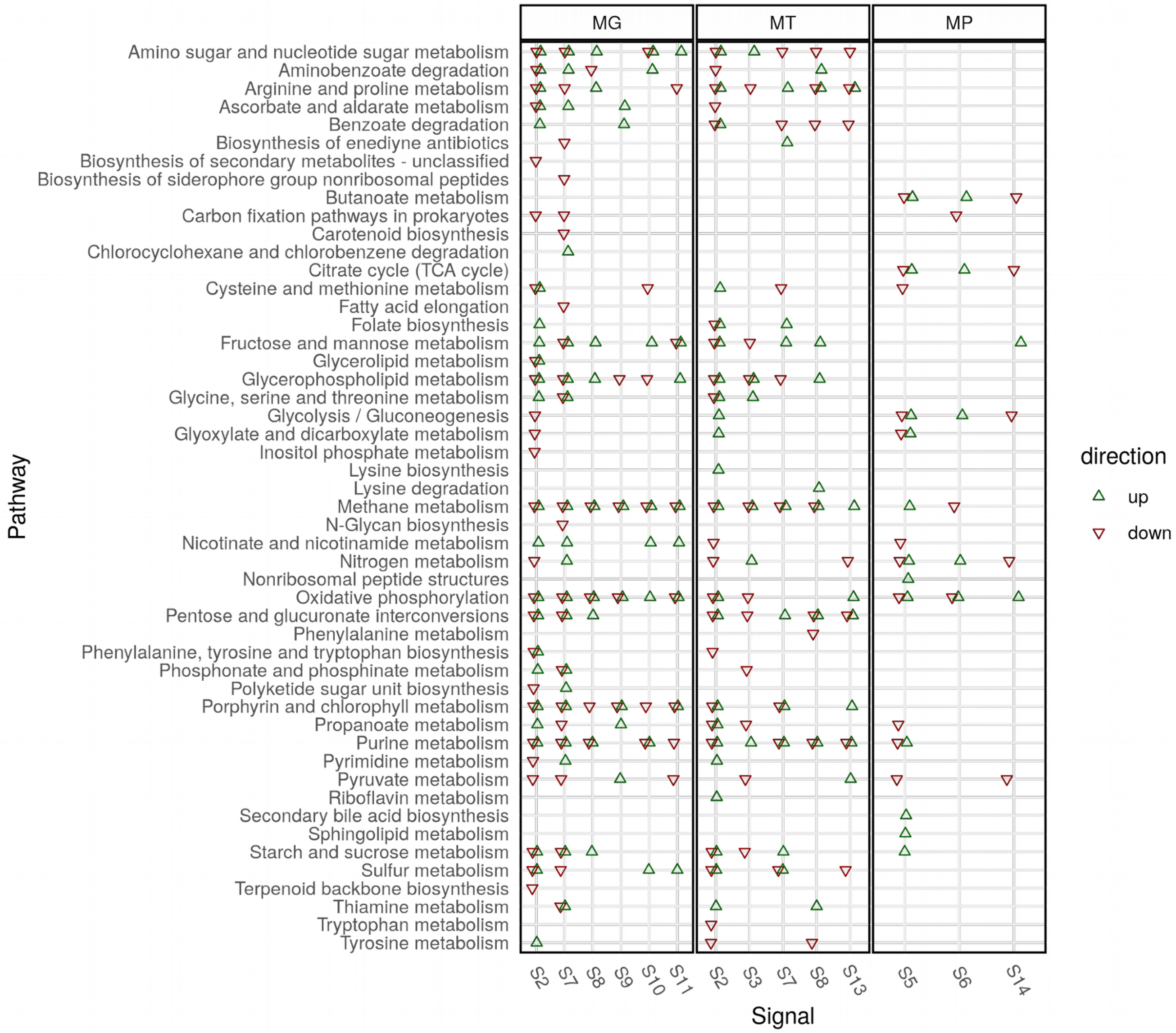


pointing up marks the intersection. Similarly if the loading is in the bottom 5% (strongly negative) a red pointing down marks the intersection. The vertical blocks separate the three omics. The plot integrates also lower metabolic labels (i.e. KO) and some might disagree in orientation, leading to pathways with both types of arrows.

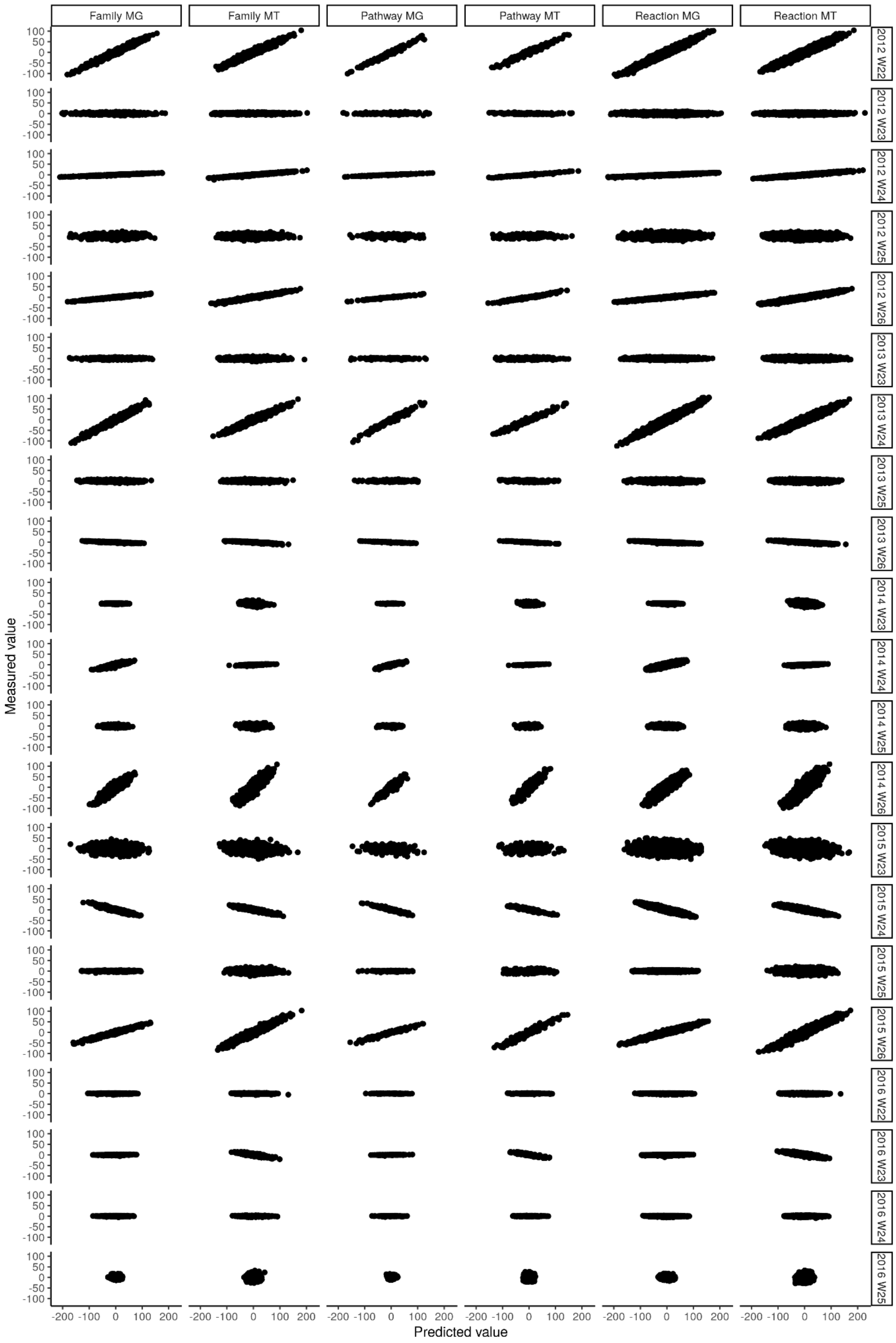


**Supplementary Figure 10**. Reconstructed abundance and gene expression of all the microbial families, reactions and pathways in the community versus the real one for each sample in the test set.

**Supplementary Table 1**. Number of representative MAGs (rMAGs), contigs (rContigs) and ORFs per biological subset (prokaryotic, eukaryotic, plasmidial and viral).

**Supplementary Table 2**. Reported p-values for the Shapiro test performed on 10 random subsets (with 5000 data points each) of the betas from the batch-effect correction of the MG ORF data. The two technical variables are the number of reads and the average read length per sample.

**Supplementary Table 3**. Reported p-values for the Shapiro test performed on 10 random subsets (with 5000 data points each) of the betas from the batch-effect correction of the MT ORF data. The two technical variables are the number of reads and the average read length per sample.

**Supplementary Table 4**. Environmental parameters manually collected by the researchers at the sampling site.

**Supplementary Table 5**. Environmental parameters automatically collected by the sensors of the WWTP.
